# Transferring active inference to newly encountered prism-shifted environments

**DOI:** 10.1101/2025.11.10.687533

**Authors:** Megumi Yoshihara, Takuya Isomura

**Affiliations:** Graduate School of Informatics, Kyoto University, 36-1 Yoshidahonmachi, Sakyo-ku, Kyoto 606-8501, Japan; Brain Intelligence Theory Unit, RIKEN Center for Brain Science, 2-1 Hirosawa, Wako, Saitama 351-0198, Japan

**Keywords:** Free-energy principle, Prism adaptation, Active inference, POMDP, Transfer learning

## Abstract

Active inference has been proposed as a unified explanation of perception and action. Previous work has shown that canonical neural networks that minimize shared Helmholtz energy are cast as performing active inference of the external environment. However, how animals flexibly adapt to newly encountered environments remains to be fully addressed. To investigate the brain’s generalizability and adaptability to new environments, this work develops canonical neural networks that employ multiple policy matrices in parallel. We demonstrate that the proposed model can recapitulate the prism adaptation—a form of visuomotor adaptation—under an arm-reaching task. Using policy matrices pretrained under various target positions, these networks could transfer previous experiences and exhibit faster adaptation to the prism-shifted environment than the naive networks. Furthermore, after-effects were observed following the removal of simulated prism glasses. These results suggest the biological plausibility and utility of the proposed model, providing insights into the adaptive capabilities of the brain.

## INTRODUCTION

The brain exhibits great adaptability to newly encountered environments, and modeling it using neural networks remains a significant challenge in neuroscience. The free-energy principle (FEP) (Friston et al., 2006; Friston, 2010) has been proposed to account for perception (Lanillos et al., 2021; Mirza et al., 2021), learning, and action(Klar et al., 2025; Lanillos et al., 2021) of biological organisms in terms of Bayesian inference and the minimization of variational free energy. Biological organisms are considered to develop the generative model—a hypothesis about the dependence structure between hidden states and sensory inputs—in the brain by minimizing variational free energy as a tractable proxy for minimizing the sensory surprise. By doing so, they perceive hidden environmental states and optimize their actions accordingly to minimize the risk associated with future outcomes, a process referred to as active inference (Da Costa et al., 2020; Friston et al., 2017).

Notably, recent works have shown that canonical neural networks that minimize Helmholtz energy can be read as following the FEP under a class of partially observable Markov decision process (POMDP) models (Isomura et al., 2022). These networks can exhibit active inference under various task settings, including causal inference (Isomura & Friston, 2020) rule learning (Tazawa & Isomura, 2024), prediction, and planning (Paul et al., 2024) These properties suggest that recapitulating the external environment in the network’s internal states is an inherent property of neural networks.

Despite these successes, when encountering novel environments, these networks basically need to learn a generative model for new environments from scratch. This requires a considerable training cost, and the generalization error usually declines with a widely observed power-low order of time (i.e., 𝒪 (*t*^−1^)) (Hastie et al., 2009). By contrast, recent advances in machine learning suggest that transferring previously learned models may facilitate these networks to adaptation to new environments (Kouw & Loog, 2018; Zhuang et al., 2020). However, whether canonical neural networks can exhibit such transfer learning remains largely unexplored.

To investigate the neuronal mechanisms of adaptability and generalizability, prism adaptation serves as a representative visuomotor phenomenon (Hermann & Cattell, 1898; Prablanc et al., 2020). This can be observed, for instance, when subjects perform a reaching task while wearing the prism glasses that shift their visual field. Initially, subjects wearing prism glasses struggle to complete reaching tasks as prediction misalignment causes an increase in reaching error; after a few trials, however, they gradually learn to accurately point to targets with minimal errors, demonstrating the adaptability to new environments. Upon removal of the prism glasses, initial errors are made but are immediately recovered. This task requires the readjustment and integration of visual and somatosensory information to novel sensory-motor contingencies, making it suited to model the brain’s generalizability using canonical neural networks.

In doing so, some observations can be considered as a generalizability criterion: In healthy human prism exposure, the adaptation occurs within approximately 10 trials, and the reaching error approaches approximately zero (Luauté et al., 2009; Rossetti et al., 1993). This is a remarkably low number compared to the number of times a person has ever reached, indicating that biological brains can immediately correct their predictive discrepancies. However, previous modeling works have yet to fully explain the neuronal mechanisms underlying this significant gap (Sakaguchi et al., 2001; Smith et al., 2006; Inoue et al., 2015). Although Bayesian frameworks have been proposed as a promising avenue for addressing this issue (Petitet et al., 2018), concrete models that capture the underlying processes are still lacking.

To address this issue, this work developed canonical neural networks that exhibit transfer learning based on the FEP and active inference and applied them to explain the prism adaptation. We demonstrated that these neural networks can transfer previously learned policy matrices— pretrained to generate actions to reach various target positions depending on observations—to facilitate adaptation to a newly encountered environment. These networks exhibited the rapid adaptation in the presence of simulated prism glasses and the after-effects following the removal of the prism glasses, as observed in empirical prism adaptation. Mathematical analyses demonstrated that the prediction error for selecting the optimal mixture of policy matrices converges with an exponential order decay (i.e., 𝒪 (*e*^−*α t*^) with a coefficient *α* > 0), which is considerably faster than the widely known *t*^−1^ order decay of generalization errors for conventional models. We conclude by discussing possible biological implementations of these learning mechanisms.

## METHODS

### Canonical neural networks

We began by outlining canonical neural networks (Isomura et al., 2022; Isomura & Friston, 2020) with a particular focus on modeling arm reaching tasks. We defined a two-layer canonical neural network (Fig. 1) of rate-coding models comprising middle 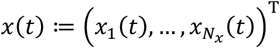 and output 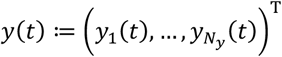 layers. On receiving sensory inputs 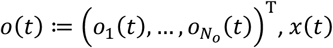 yields the neural dynamics and propagates it to *y*(*t*); then, *y*(*t*) generates actions of the agent, that is, a motion in either up, down, left, or right direction in a grid world. These neural activities are provided as follows:

**Fig. 1.**
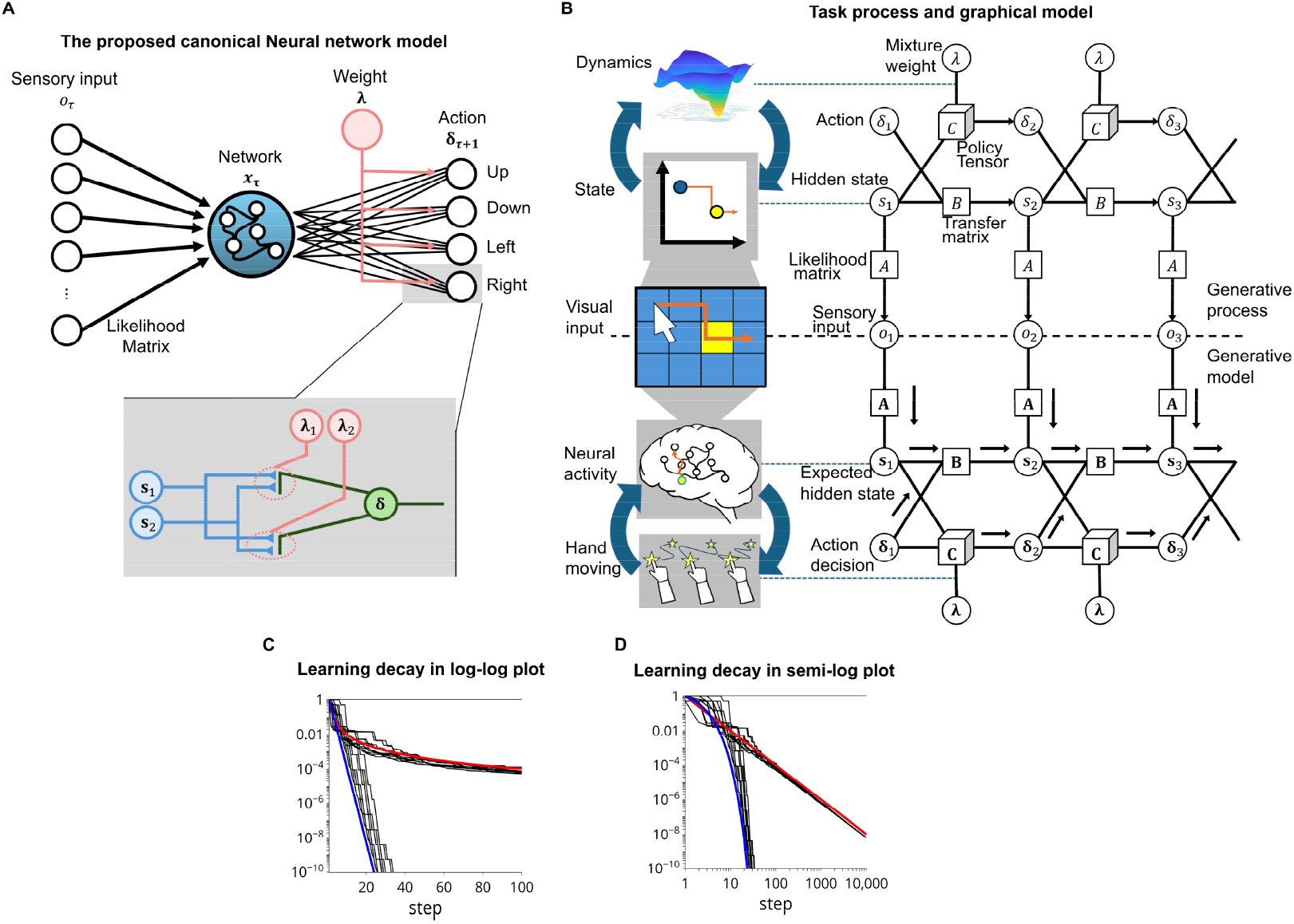
Schematic of canonical neural networks performing active inference. **A**. Canonical neural network architecture for performing an arm reaching task (top) and possible neuronal implementation of policy learning in terms of the three-factor learning rule (bottom). Middle layer activity *x*_*τ*_ is generated by sensory inputs, 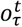 and 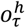 that carry information about target and hand positions. Action ***δ***_*τ*+1_ is generated by *x*_*τ*_ using the tensor of policy matrices, yielding a hand motion in the grid world. **B**. A POMDP model for reaching tasks expressed in the form of a graphical model. The lower part formally corresponds to the canonical neural network shown in (A). This architecture enables transfer learning by selecting *C* from the pool via attentional switch *λ*. Network weights and *λ* are updated by using *s*, ***δ***, and risk *Γ*. **C**. Generalization errors in **C** and *λ*. **D**. Engaged view of (C) with semi-log scale.

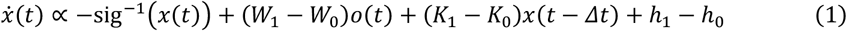

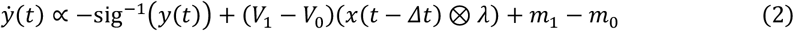

where 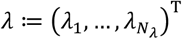 denotes firing intensities of modulator neurons encoding mixture parameters; 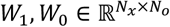 and 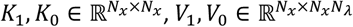 are synaptic weight matrices; and *h*_1_ ≔ *h*_1_(*W*_1_, *K*_1_), *h*_0_ ≔ *h*_0_(*W*_0_, *K*_0_), *m*_1_ = *m*_1_(*V*_1_), *m*_0_ = *m*_0_(*V*_0_) are the adaptive firing thresholds that depend on synaptic strengths. Here, (*V*_1_ − *V*_0_)(*x*(*t* − *Δ t*) ⊗ *λ*) represents synaptic inputs from the middle layer that are modulated by *λ*.

Without loss of generality, Equations (1) and (2) can be derived as a gradient descent on Helmholtz energy 𝒜. The functional form of 𝒜 can be identified by computing the integral of the right-hand side of Equations (1) and (2), as follows:

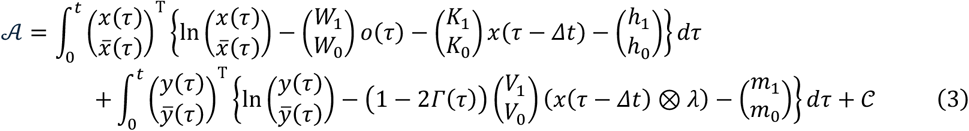

The form of 𝒜 is equivalent to the variational free energy under a class of POMDPs (Isomura et al., 2022). This indicates that the dynamics of canonical neural networks are equivalent to Bayesian Belief updating under a class of POMDPs, supporting the validity of the free energy principle as a universal characterization of neural dynamics and self-organization. Notably, a gradient descent on the shared Helmholtz energy derives the synaptic plasticity in the output-layer synaptic weights *V*, which is expressed as neuromodulation of Hebbian plasticity in the form of a three-factor learning rule (Frémaux & Gerstner, 2016; Kuśmierz et al., 2017; Pawlak et al., 2010) (Fig. 1A). This formally corresponds to the update of policy matrix to minimize future risks (Isomura et al., 2022). Based on this foundation, hereafter we formulated prism adaptation in terms of POMDPs (that correspond to canonical neural networks) following the FEP.

### Corresponding generative models

The behavior of an agent interacting with a discrete state space is expressed in the form of a POMDP as follows: The environment comprises binary sensory inputs (observations) 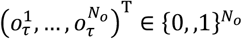 and hidden states 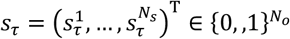, and their sequences are denoted as *o*_1: *t*_ = {*o*_1_, …, *o*_*t*_} and *s*_1: *t*_ = {*s*_1_, …, *s*_*t*_}. Here, hidden states involve a factorial structure comprising two one-hot vectors of hand 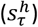 and target 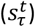 positions, which is denoted as 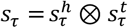 using the Kronecker product operator ⊗. Similarly, sensory inputs 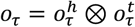 comprise a product of somatosensory input for hand position 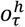 and visual input for target position 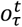. Action or decision *δ*_*τ*_ describes the motion of the agent’s hand, which can move up, down, left, or right at each time step. This makes the agent’s hand position (*s*^*h*^) move one grid, but it will stay if it reaches the end of the grid field.

Sensory inputs *o*_*τ*_ are generated based on hidden states *s*_*τ*_ in the form of a categorical distribution *P*(*o*_*τ*_|*s*_*τ*_, *A*) = Cat(*As*_*τ*_) through the likelihood matrix *A*. The dynamics of hidden states depend on the previous state and action, expressed as *P*(*s*_*τ*+1_|*s*_*τ*_, *δ*_*τ*_, *B*) = Cat(*B*(*δ*_*τ*_ ⊗ *s*_*τ*_)) using transition matrix *B*. The agent’s action is determined based on the policy matrix *C* that defines the selection probability of action depending on hidden states, *P*(*δ*_*τ*+1_|*s*_*τ*_, *C*) = Cat(*Cs*_*τ*_).

Here, a counterfactual learning of the policy matrix is adopted following the previous work (Isomura et al., 2022). Under this setting, the agent adopts a risk-dependent policy model 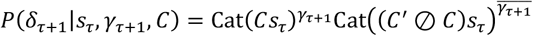 to update *C*, in which the agent learns to retain or enhance the current strategy if the risk *γ*_*τ*+1_ is 0, whereas the agent forgets it if the risk is 1 (where 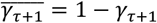 denotes the sign-flipped *γ*_*τ*+1_). Hence, the generative model is defined as follows:

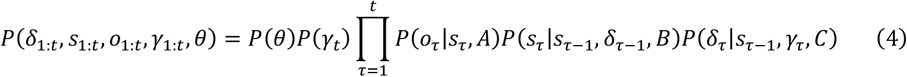

where *θ* is a set of parameters, *θ* = {*A, B, C*}. Prior beliefs about parameter matrices *P*(*θ*) = *P*(*A*)*P*(*B*)*P*(*C*) follow Dirichlet distributions *P*(*A*) = Dir(*a*), *Q*(*B*) = Dir(*b*), and *Q*(*C*) = Dir(*c*) parameterized by Dirichlet parameters *a, b*, and *c*.

### Implementation of transfer learning

In contrast to naive active inference agents defined above (i.e., naive learners), here we define agent exhibiting transfer learning (i.e., transfer learners). Transfer learning (Zhuang et al., 2020) enables the deployment of previously learned policy matrices to quickly adapt to newly encountered environments, including the presence of prism exposure. In this work, we model policy matrix *C* using a mixture of policy matrices as a function of mixture parameter *λ*, as follows:

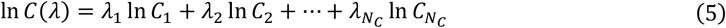

Each fixed independent policy matrix *C*_*k*_ defines the selection probability of hand action in the *k*-th context, and *λ* is a mixture weight parameter or attentional switch that follows a categorical distribution *λ*~Cat(***Λ***), analogous to a mixture of generative models(Isomura et al., 2019). A set of parameter matrices {*C*_*k*_} are generated by pretraining using naive learners. In this formulation, transfers learners employ the parameter set *θ* = {*A, B, λ*} and update them instead of updating *θ*_naive_ = {*A, B, C*}.

### Variational Bayesian inference

An approximate posterior distribution is defined using a mean-field approximation as

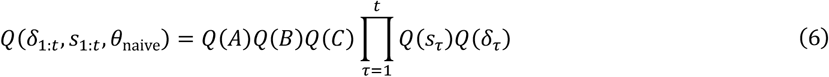

for naive learners and

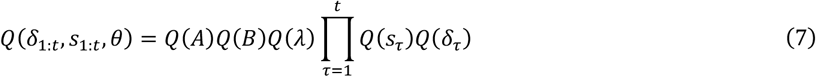

for transfer learners. Approximate posterior beliefs about hidden states, action, and mixture parameter follow categorical distributions *Q*(*s*_*τ*_) = Cat(*s*_*τ*_), *Q*(*δ*_*τ*_) = Cat(***δ***_*τ*_) and *Q*(*λ*) = Cat(*λ*) parameterized by the posterior expectations *s*_*τ*_, ***δ***_*τ*_, and *λ*. Moreover, approximate posterior beliefs about parameter matrices follow Dirichlet distributions *Q*(*A*) = Dir(*a*), *Q*(*B*) = Dir(*b*) and *Q*(*C*) = Dir(*c*) parameterized by Dirichlet parameters *a, b*, and *c*.

Given the above generative model and posterior distribution, the variational free energy for naive learners is provided as follows:

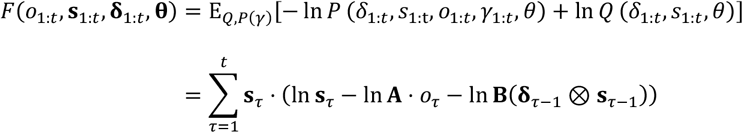

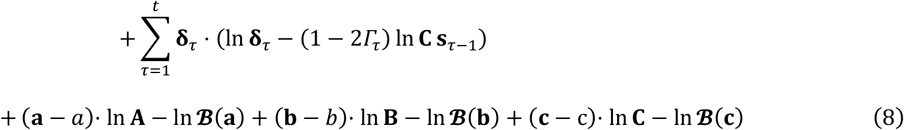

For transfer learners, the complexity of *C* in Equation (8), (*c* − *c*)· ln **C** − ln **ℬ**(*c*), is replaced with that of *λ*, which is given as 𝒟_KL_[*Q*(*λ*)||*P*(*λ*)] = *λ* · (ln *λ* − ln *Λ*). Here, *Γ*_*t*_ ∈ [0,1] is a risk that evaluates the goodness of action for each step.

The gradient descent on *F* furnishes the Bayesian belief update rules. Posterior beliefs about hidden states *s*_*τ*_ and action *δ*_*τ*_ are updated for each step. These inference update rules are derived by solving the fixed point of implicit gradient descent (i.e., *∂F*/*∂s*_*t*_ = 0 and *∂F*/*∂****δ***_*t*_ = 0):

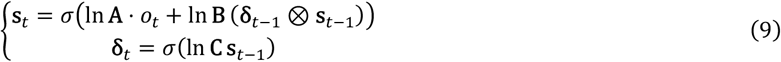

Dirichlet parameters *a, b, c*, and mixture parameter *λ* are updated at the end of each trial as part of the learning. These learning update rules are derived by solving *∂F*/*∂θ* = 0 as:

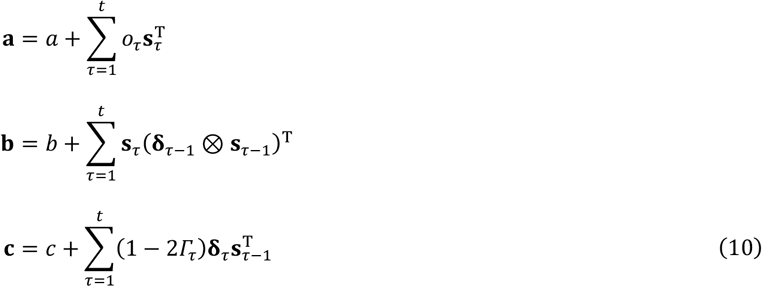

for naive learners. For transfer learners, the *C*’s update is replaced with the *λ*’s update as:

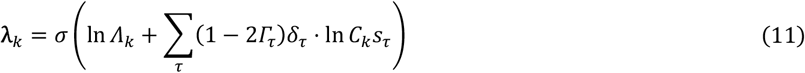

Through Equation (11), the agent learns to select the action that minimizes the risk, while avoiding actions that increase the risk.

### Simulation settings

In the simulation, a set of 100 policy matrices, 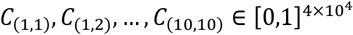, are prepared for a mutually different target position, where each *C*_(*n,m*)_ denotes the one pretrained to reach the target on the coordinate (*n, m*). During each pretraining, the target position was fixed at (*n, m*), and the agent continued to randomly move its hand for 100000 steps, which was followed by the learning of the posterior belief about *C*_(*n,m*)_ based on Equation (10). In pre-training, we use an *A* matrix created based on a uniform distribution as the initial value so that the same policy can be adopted even if the target moves to a nearby location.

## RESULTS

### Transferring active inference to novel environments

This work aims to mathematically model the generalizability and adaptability of the brain to novel environments. An arm reaching task is a widely used visuo-motor paradigm to investigate the emergence of sensory-motor contingencies, in which subjects move their hands towards the target position based on visual input (Bourgeois et al., 2021). This can be modeled using canonical neural networks (Isomura et al., 2022), which receive visual and somatosensory inputs, infer the latent environmental states via middle-layer neural activity, and generate actions to shift the arm position (Fig. 1A, top). Risk here is characterized by the distance from the target, and the network learns the optimal action policy through the plasticity of the output layer synaptic connection weights (Fig. 1A, bottom). As previously established (Isomura et al., 2022), canonical neural networks are read as performing active inference of the external environment to minimize risks associated with future outcomes, under a POMDP generative model (Fig. 1B). Thus, after sufficient exposure to the environment, these networks self-organize to recapitulate the interacting environment as a universal property.

In doing so, conventional models need to relearn the policy matrix (*C*) from scratch upon encountering a new environment, making it extremely time-consuming. Since the policy matrix posterior takes continuous values following a Dirichlet distribution, this method exhibits a widely observed 1/t-order generalization error decay (Ref: Hastie et al., 2009) (Fig. 1C, D, red lines). However, this indicates a large gap from humans and animals that can quickly adapt to moving their arms to a given target position (Luauté et al., 2009; Rossetti et al., 1993).

To model the quick adaptation, we constructed canonical neural networks that employ multiple policy matrices in parallel, each of which was pretrained to reach different target positions. The key advantage is the incorporation of the switching parameter λ that follows a categorical distribution. In contrast to the naive model, this network converges the generalization error considerably faster in a simple example setting (Fig. 1C, D blue lines).

Mathematical analyses revealed the distinct learning mechanisms (Appendix A). Critically, discrete switching variable *λ* tends to converge to a one-hot vector as evidence is accumulated. This makes the optimal policy selection from a set of policies converge with an exponential order, a speed considerably faster than the well-known 1/t order generalization error decay. Intuitively, this is because upon receiving sensory inputs, the evidence is accumulated linearly with time (i.e., 𝒪 (*t*))—and the amplitude of the selected value is determined by the difference in evidence for each component of *λ*. This makes the posterior expectation in the form of an exponential function exp(−*t*⟨evidence gap⟩), where ⟨evidence gap⟩ denotes the difference in normalized evidence. Although computing this difference involves the generalization error in the 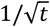 order, this is negligible compared to the order 1 amplitude of the difference itself. Therefore, the generalization error of *λ* is determined by the form exp(−*t*(*X*_1st_ − *X*_2nd_)) as the leading order, where *X*_1*s t*_ and *X*_2*nd*_ are the normalized evidences for the most plausible and second most plausible policies, causing it to decrease exponentially with time. Further details are provided in Appendix A.

In essence, both mathematical analyses and numerical simulations (Fig. 1C, D) suggest that the selection mechanism of policy matrices enables more rapid adaptation than naive parameter learning, analogous to empirically observed prism adaptation. Based on these observations, in the remainder of this paper, we formally model prism adaptation using the canonical neural networks exhibiting transfer learning—and examine whether the rapid adaptation to a prism-shifted environment and the after-effects following the removal of prism glasses can be reproduced.

### Modeling prism adaptation

In this section, we instantiated transfer of active inference through modeling a prism-shift paradigm of arm reaching tasks (Hermann & Cattell, 1898; Rossetti et al., 1993) (Fig. 2A). Agents perform a reaching task. For each trial, the agent’s hand is initially located at a random position in a 10 × 10 grid field (Fig. 2B). At each step, the agents receive sensory information about target and hand position and infer hidden states and actions to determine the next motion from either up, down, left, or right (Equation (9)). A trial ends when either hand reaches the target or 100 steps have passed, followed by the updates of parameters (Equations. (10) and (11)). This cycle continues for 200 trials. The risk *Γ*_*t*_ is defined based on the change in Manhattan distance between the hand and target: *Γ*_*t*_ = 0.45 when the distance decreases for each step (i.e., when the hand moved toward the target); otherwise, *Γ*_*t*_ = 0.55.

**Fig. 2.**
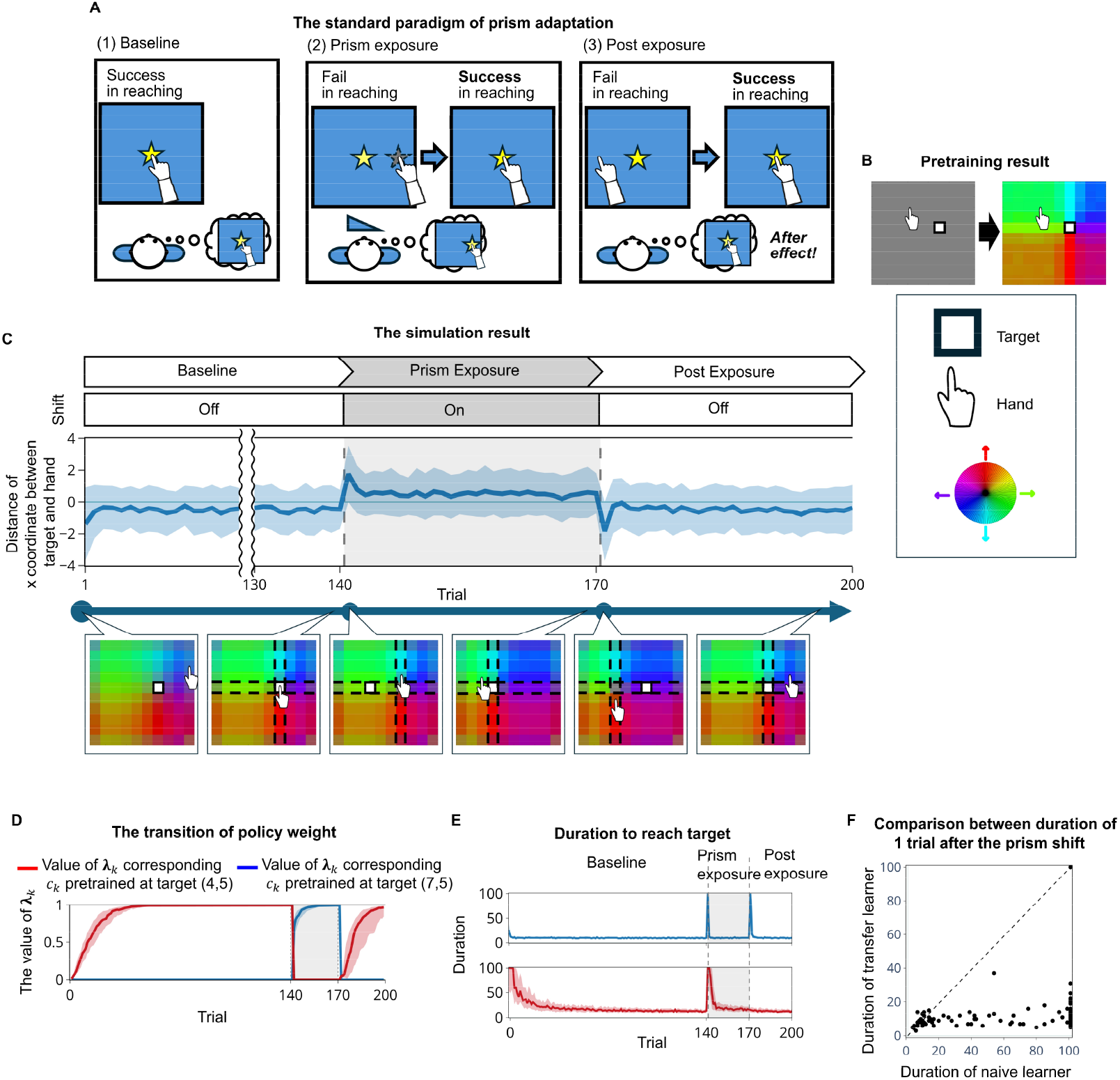
Simulating prism adaptation using canonical neural networks. **A**. The prism adaptation paradigm: (1) a subject performs an arm reaching task in the baseline phase; (2) during the prism exposure phase, the same task is performed with wearing prism glasses; and (3) during the post-exposure phase, the prism glasses are removed. **B**. The pretraining of policy matrices. Prism adaptation is modeled with a discrete state space (POMDP) model. The hand and target are placed in a 10×10 grid world. The colors in each grid represent the hand-moving directions that correspond to the color map besides the heatmap, indicating the action selection probability for moving up/down/left/right. **C**. Overview of the simulation setting. The upper panel shows the trajectory of the x-coordinate difference between the target and hand after the minimum steps. The bottom panel shows the corresponding changes in the policy matrix. Heatmaps illustrate how the agent’s policy matrix changes over trials. While the colors are blurred before training (leftmost), the colors become high contrast through training. The fixed point (crosspoint of vertical and horizontal dashed lines) approximately pursues the target position (open square) both in the absence and presence of prism shift. **D**. Changes in the two policy mixture weights corresponding to original and new target positions. **E**. The upper and lower panels show the duration (the number of steps) for transfer learners and naive learners, respectively. **F**. Comparison of the duration at the 1 trial after prism exposure between the transfer and naive learners. In (C)–(E), the lines and shaded areas indicate the medians and interquartile ranges.

A simulation comprises three phases (Fig. 2C): The first 140 trials serves a “baseline” environment without prism glasses, in which the target is placed on the coordinate (*x, y*) = (7,5) (Fig. 2C, Baseline). Starting from a flat prior *λ*, the agent optimized the mixture parameter *λ* for the baseline condition (Fig. 2C). The second 30 trials correspond to the “prism exposure,” in which the target is placed on (*x, y*) =(4,5) (a white filled grid in Fig. 2C, prism exposure phase) but its apparent position is still observed at (7,5). This prism shift was modeled by changing the likelihood mapping *A*, which displaces only the target position but not the hand position. While the apparent target position is unchanged from the baseline, the agent can notice the difference when it receives the risk after an action was made. In the last 30-trial “post exposure” phase, the agent returns to the environment in the absence of prism exposure, where the target is placed—and observed—at (*x, y*) = (7,5). In each phase, the agent learns a policy that brings the hand closer to the true target by minimizing the free energy.

We demonstrated that transfer learners that employ pretrained policy matrices can replicate the key empirical properties of prism adaptation. Performance was assessed by the error between the x coordinates of the target and hand after the minimum steps to reach the target (Fig. 2C). While large errors were initially observed during both the prism exposure and post-exposure phases, they rapidly diminished to approximately zero. The transition in policy matrix **C** shows the occurrence of adaptation in both phases (Fig. 2C, bottom), which was driven by the update of mixture weights *λ* (Fig. 2D). In each phase, transfer learners consistently learn the optimal *λ* values. Specifically, during prism exposure, *λ* is adjusted to align the policy with the actual target position, while in the post-exposure phase, *λ* reverts to the baseline policy.

We observed distinct properties between transfer and naive learners. The proposed model (transfer learner) qualitatively replicated experimental outcomes (Fig. 2E). During prism exposure, initially large duration to reach the target decreased after approximately 5 trials and converged around 10. In the post-exposure phase, transfer learners initially exhibited an after-effect akin to empirical prism adaptation and re-learn gradually. In contrast, naive learners that learn policy matrix **C** directly via Equation (10) could not reproduce the after-effect (Fig. 2E, red error bars). While the duration of naive learners increased during the prism exposure, they failed to exhibit the duration increase during post-exposure, arguably because of slow learning during the prism exposure.

In summary, the proposed model can learn to perform the arm reaching task both in the absence and presence of prism shift in the Bayes optimal manner and replicate the dynamics of empirical prism adaptation. We confirmed that these outcomes can be observed even with randomly varied target positions (Fig. 3).

**Fig. 3.**
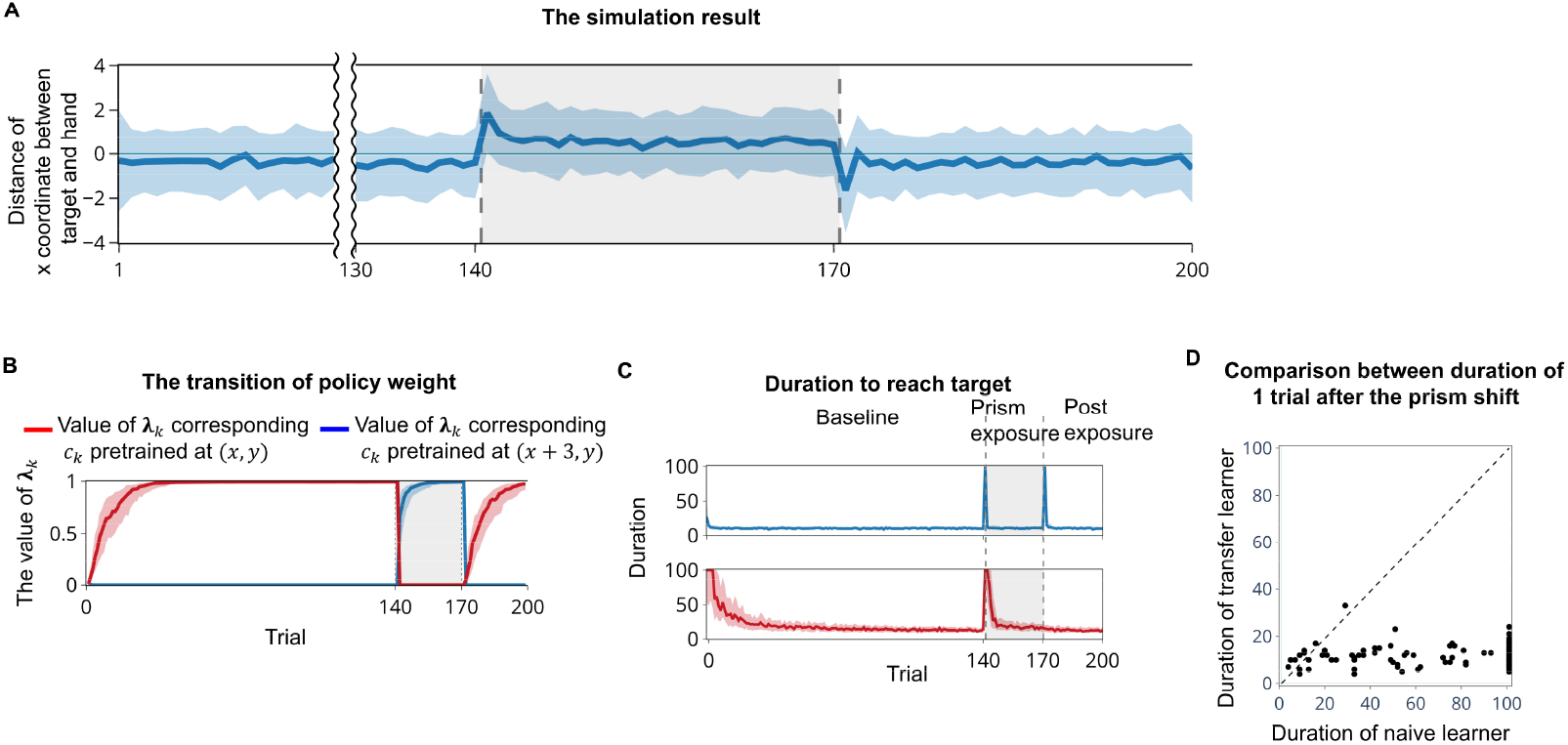
Simulations with randomized target. Same as Fig.2 but the target position is changed randomly over sessions, showing the robustness of the proposed model. **A**. The x-coordinate difference between the target and the hand after the minimum steps. **B**. Changes in the two policy mixture weights. **C**. Duration for the transfer (top) and naive learners (bottom). **D**. Comparison of the duration value between the transfer and naïve learners. In (A)–(C), the lines and shaded areas indicate the medians and interquartile ranges.

### Exponential-order generalization error decay in prism adaptation

Lastly, we analytically investigated the learning speed of transfer and naive learners. For analytical tractability, here we assume that the posterior hidden state *s*_*τ*_ well approximates the true hidden state *s*_*τ*_, the action *δ*_*τ*_ follows a random walk, and that there exists an optimal policy matrix *C*^*^ in a set of pretrained matrices {*C*_*k*_}. Under these conditions, the mixture parameter *λ* and policy matrix **C**(*λ*) converge to the Bayes optimal values *λ*^*^ and **C**^*^ with the *e*^−*a t*^ order decay of generalization error, where *a* > 0. Detailed derivations are described in detail in the Appendix. This convergence speed is considerably distinct from learning speed in the policy matrix **C** of naive learners that converges to the Bayes optimal value **C**^*^ with the 1/*t* order decay.

To valid the change in generalization error, we simulated the transfer learners under the condition in which the action is defined as a random walk and confirmed the *e*^−*a t*^ order convergence of policy mixture weights *λ* (Fig. 4A) and policy matrix **C** (Fig. 4B). The generalization error of the mixture weight *λ* in transfer learners converges in approximately 2400 steps, and its change resembles a straight line in a semi-log plot, indicating the exponential order convergence (Fig. 4A, blue line). After 2400 steps, more than half of *λ* elements become Bayes optimal one-hot vector; therefore, the median generalization error becomes zero. The gradient of the generalization error in the semi-log plot closely matches the theoretical gradient calculated in the Appendix. The generalization error of the policy matrix **C** in the transfer learner also exhibited a similar property (Fig. 4B), in which empirical errors resembled a theoretical line (green line). In contrast, the naive learner requires a longer convergence time in the *t*^−1^ order (Fig. 4C), aligning a line in a log-log scale after about 500 steps.

**Fig. 4.**
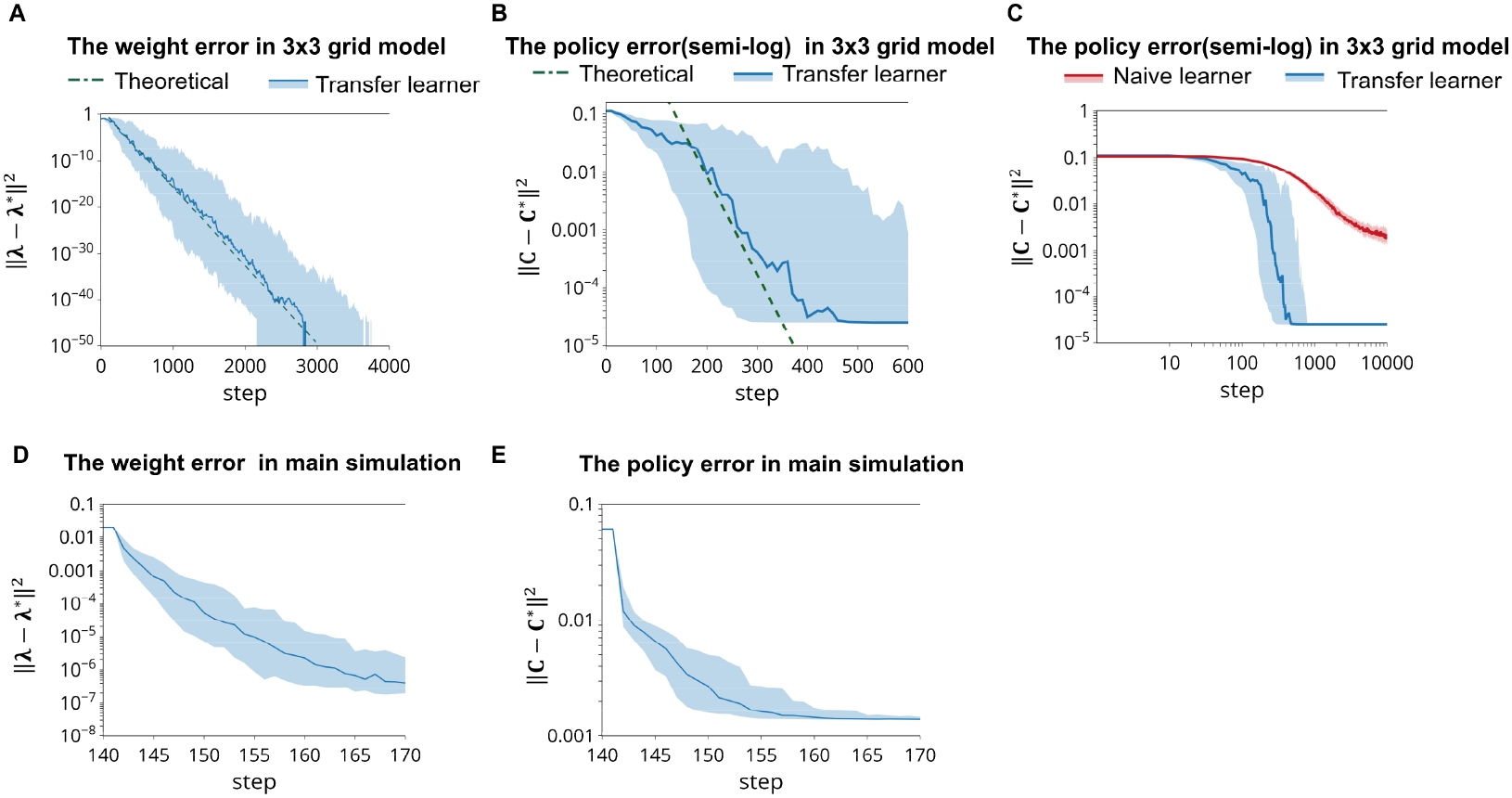
Generalization error in simplified model simulation. **A**. Generalization error of *λ* and the theoretical value in toy model. **B**. Generalization error of **C** and theoretical value in semi-log plot in toy model. **C**. Generalization error of **C**, and theoretical value in log-log plot. Both learners have a common initial value of the policy matrix in toy model. **D**. Generalization error of *λ* in main simulation. **E**. Generalization error of **C** and theoretical value in semi-log plot in main simulation.

In essence, the error of the transfer learner decreases much faster than that of the naive learner, enabling a faster optimization of the policy matrix. The faster adaptation is further confirmed under the condition where the hidden states are updated following Bayesian belief updating rules (Fig. 4D, E). At the one trial after the prism shift (i.e., trial 142), the majority of transfer learners reached the target significantly faster than the naïve learners (Fig. 2F; *p* = 4.15 × 10^−18^, Mann–Whitney U test) and the median duration of transfer learners was only approximately 8.3% of that of naive learners.

## DISCUSSION

In this work, we developed an active inference model incorporating transfer learning and applied it to model prism adaptation. Canonical neural networks employing multiple policy matrices can transfer previously learned policies to novel environments, demonstrating faster adaptation than naive learners. This has broader implications for understanding the brain’s generalizability and adaptability, particularly in the context of prism adaptation. The transfer learners involve three different timescales comprising fast updates of state *s* and action λ, slow updates of mixture weights *λ*, and ultra-slow updates of parameters **A, B** and **C**, akin to previous computational models for prism adaptation (Inoue et al., 2015).

Two timescales in prism adaptation were suggested by several works (G. M. Redding et al., 2005; G. Redding & Wallace, 2002, 2006). On this view, fast and slow adaptations are interpreted as recalibration of frames and realignment of the visuo-motor mapping, respectively, and the latter induces after effect. Our observations were consistent with this view: only transfer learners that update *λ*, but not naive learners without *λ*, exhibited after effect, indicating the importance of transfer mechanisms to reproduce empirical phenomena.

Such correspondences in timescales are beneficial to identify a precise mapping between the parameters of active inference models and corresponding brain regions. Specifically, faster timescale adaptation is attenuated by lesions in the posterior parietal cortex (Canavan et al., 1990; Clower et al., 1996; Newport & Jackson, 2006; Pisella et al., 2004; Welch & Goldstein, 1972). This impairs both error reduction and after effects. Moreover, fast adaptation involves the primary motor cortex (Danckert et al., 2008), anterior cingulate cortex (Danckert et al., 2008; Luauté et al., 2009), and cerebellum (Danckert et al., 2008; Luauté et al., 2009). In particular, cerebellar lesions result in decreased or absent prism after-effects (Calzolari et al., 2015; Hanajima et al., 2015; Pisella et al., 2005), suggesting its role in slow adaptation.

Furthermore, the underlying neuronal mechanisms can be considered in terms of neuromodulation of synaptic plasticity. The *λ* update rule in Equation (11) compares action and experience and reinforces optimal policy, providing a functional explanation of policy adaptation, which may be associated with attentional switch and working memory mediated by such as dopaminergic neurons. Then, the process of amplifying more similar connections and attenuating less similar connections can be formulated as dopaminergic modulation of synaptic plasticity.

Many traditional models in prism adaptation have posited multi-timescale mechanisms (Inoue et al., 2015; Kim et al., 2015; Kording et al., 2007). While these models can account for behavioral performance, they do not fully explain the underlying neuronal mechanisms and how different timescales contribute to learning. The proposed canonical neural networks have potential to assimilate empirical data into modelling. Earlier work has established a reverse engineering method to estimate generative model from empirical neural activity data based on canonical neural networks (Isomura et al., 2023, 2025). This allows us to compare model predictions with empirical results (Kitazawa et al., 1995). In future work, we hope to elucidate the functional relationship between brain regions and the prism adaptation process by comparing the network properties of our model and empirical data.

## CONCLUSION

In summary, we developed a transfer learning model of active inference based on canonical neural networks. Our model selected the Bayes optimal parameter from pretrained parameters and demonstrated faster adaptation to a novel prism glasses environment. Further, it successfully recapitulated features of prism adaptation such as after effect. These results support that the mixture policy matrices enable transfer of knowledge learned from experiences to quickly adapt to the environment and have the capacity to reproduce the brain’s generalizability.

## Appendix Comparison of generalization errors

Here, we assume that the posterior beliefs about the hidden state and action well approximate the true hidden state and action: *s* = *s*, ***δ*** = *δ*. In addition, we assume that {*δ*_*τ*_} is a random walk, indicating that any 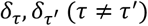 are independent and identically distributed.

We define a Hebbian product as a block vector, 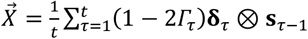 and its large-time limit as 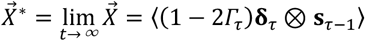. The generalization error of 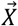 is given by the covariance matrix of their difference 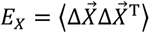, where 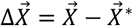. When the agent obeys a random work, *E*_*X*_ can be analytically computed, showing the *t*^−1^ order convergence.

The posterior belief about the policy mapping *C* is parameterized by the Dirichlet parameter, which is given as Equation (7), or equivalently, as a vector 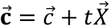. Then, the posterior belief about *C* is given as 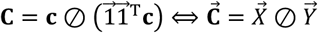 when *c* is negligibly small, where ⊘ indicates the Hadamard division operator and 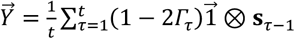. When the agent obeys a random walk, 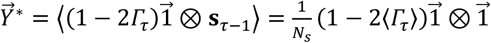 holds. This provides 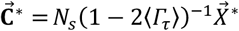.

The generalization error of 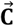 is given as 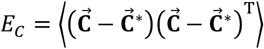. Then, 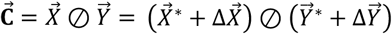 is approximated as 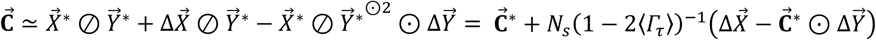 as the first-order approximation. Thus, *E*_*C*_ becomes

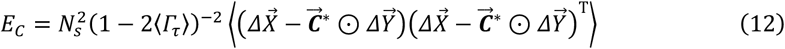

Because 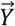 is expressed as 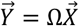 using a block diagonal matrix 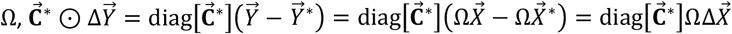 holds. Thus, we obtain

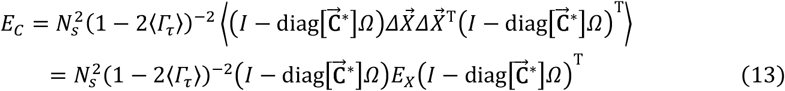

In contrast, the transfer learner that equips with the mixture policies updates the mixing balance *λ* instead of updating *c* directly. In this case, the posterior belief about the policy mapping is given as follows:

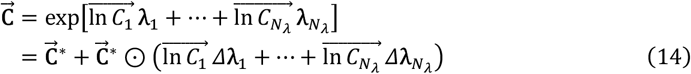

Here, the *λ* update rule is expressed as 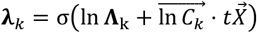 and its large-time limit as 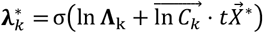. The difference between 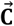 and 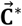 is computed as

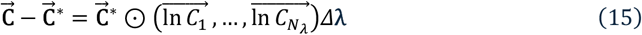

where Δ*λ* = *λ* − *λ*^*^ holds and *Λ* is assumed to be a flat prior and thus omitted for simplicity.

When one of the previously experienced contexts has a considerably higher similarity to the current environment compared to other contexts, an element of *λ* that corresponds to the context monotonically converges to one. Under this condition, 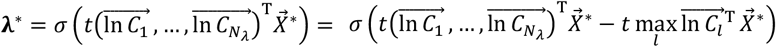 holds owing to the property of softmax function *σ*.

Thus, by defining 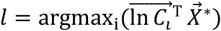 and 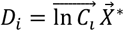, we obtain 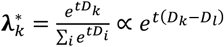. All elements of *λ* except for the largest element (*k* = *l*) converge to zero exponentially with time, i.e., 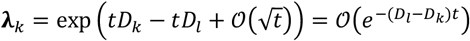 as the leading order, because the generalization error of 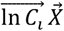 is of order 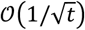 and negligibly smaller than (*D*_*l*_ − *D*_*k*_). Conversely, the largest element is computed as 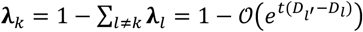, where 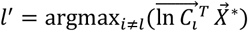 indices the second largest element.

This provides 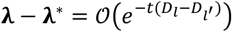. Therefore, the generalization error 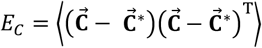 for the transfer learner is in 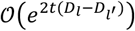, which indicates that *E*_*C*_ of the transfer learner converges to zero much faster than does that of the naive learner. Thus, the generalization error of the transfer learner is determined based on the generalization error of pretrained policy matrices.

---

**Table 1.** Glossary of expressions.

| Expression | Definition | Explanation |
| --- | --- | --- |
| $t$ | Current time | $t \in \{0, 1, \dots\}$ |
| $\tau$ | Time index | $\tau \in \{0, 1, \dots, t\}$ |
| $o_\tau$ | Sensory input | $o_\tau \in \{0, 1\}^{10000 \times 1}$ |
| $s_\tau$ | Hidden state | $s_\tau = s_\tau^h \otimes s_\tau^t$ |
| $s_\tau^h$ | Hidden state of hand position | $s_\tau^h \in \{0, 1\}^{100 \times 1}$ |
| $s_\tau^t$ | Hidden state of target position | $s_\tau^t \in \{0, 1\}^{100 \times 1}$ |
| $\mathbf{s}_\tau$ | Expected hidden state | Posterior belief of hidden state $s_\tau$,<br>$\mathbf{s}_\tau \in [0, 1]^{10000 \times 1}$ |
| $\delta_\tau$ | Action (up down left right) | $\delta_\tau \in \{0, 1\}^{4 \times 1}$ |
| $\boldsymbol{\delta}_\tau$ | Expected action | Posterior belief of action $\delta_\tau$,<br>$\boldsymbol{\delta}_\tau \in [0, 1]^{4 \times 1}$ |
| $A, B, C$ | parameter matrices | $A \in [0,1]^{10000 \times 10000}, B \in [0,1]^{10000 \times 40000},$<br>$C \in [0,1]^{4 \times 10000}$ |
| $\mathbf{A}, \mathbf{B}, \mathbf{C}$ | Expected parameter matrices | Posterior belief of parameter matrices, $A, B, C,$<br>$\mathbf{A} \in [0,1]^{10000 \times 10000}, \mathbf{B} \in [0,1]^{10000 \times 40000},$<br>$\mathbf{C} \in [0,1]^{4 \times 10000}$ |
| $\lambda$ | Mixture weight | Mixture weight for policy. |
| $\boldsymbol{\lambda}$ | Expected mixture weight | Posterior belief of mixture parameter $\lambda$. |
| $\theta$ | parameters | The set of parameters. |
| $\boldsymbol{\theta}$ | Expected parameters | Posterior belief of parameter $\theta$. |
| $a, b, c$ | Dirichlet parameter | Dirichlet parameter of matrices, $A, B, C,$<br>$a \in \mathbb{R}_{>0}^{10000 \times 10000}, b \in \mathbb{R}_{>0}^{10000 \times 40000},$<br>$c \in \mathbb{R}_{>0}^{4 \times 10000}$ |
| $\mathbf{a}, \mathbf{b}, \mathbf{c}$ | Dirichlet parameter | Dirichlet parameter of matrices, $\mathbf{A}, \mathbf{B}, \mathbf{C},$<br>$\mathbf{a} \in \mathbb{R}_{>0}^{10000 \times 10000}, \mathbf{b} \in \mathbb{R}_{>0}^{10000 \times 40000},$<br>$\mathbf{c} \in \mathbb{R}_{>0}^{4 \times 10000}$ |
| $\Gamma$ | risk | $\Gamma \in [0,1]$ |
| $\mathcal{B}(\cdot)$ | Beta function | $\mathcal{B}(\mathbf{a}_{\cdot k}) = \frac{\prod_i \Gamma(\mathbf{a}_{ik})}{\Gamma(\sum_i \mathbf{a}_{ik})}, \Gamma(\cdot)$: Gamma function |

## Acknowledgements

T.I. is supported by the Japan Society for the Promotion of Science (JSPS) KAKENHI under Grant Number JP23H04973, the Japan Science and Technology Agency (JST) CREST under Grant Number JPMJCR22P1, and the Japan Agency for Medical Research and Development (AMED) under Grant Number JP23wm0625001. The funders had no role in study design, data collection and analysis, decision to publish, or preparation of the manuscript.

